# Phylogenetic proximity drives temporal succession of marine giant viruses in a five-year metagenomic time-series

**DOI:** 10.1101/2024.08.12.607631

**Authors:** Sarah M. Laperriere, Benjamin Minch, JL Weissman, Shengwei Hou, Yi-Chun Yeh, J. Cesar Ignacio-Espinoza, Nathan A. Ahlgren, Mohammad Moniruzzaman, Jed A. Fuhrman

## Abstract

Nucleocytoplasmic Large DNA Viruses (NCLDVs, also called giant viruses) are widespread in marine systems and infect a broad range of microbial eukaryotes (protists). Recent biogeographic work has provided global snapshots of NCLDV diversity and community composition across the world’s oceans, yet little information exists about the guiding ‘rules’ underpinning their community dynamics over time. We leveraged a five-year monthly metagenomic time-series to quantify the community composition of NCLDVs off the coast of Southern California and characterize these populations’ temporal dynamics. NCLDVs were dominated by Algavirales (Phycodnaviruses, 59%) and Imitervirales (Mimiviruses, 36%). We identified clusters of NCLDVs with distinct classes of seasonal and non-seasonal temporal dynamics. Overall, NCLDV population abundances were often highly dynamic with a strong seasonal signal. The Imitervirales group had highest relative abundance in the more oligotrophic late summer and fall, while Algavirales did so in winter. Generally, closely related strains had similar temporal dynamics, suggesting that evolutionary history is a key driver of the temporal niche of marine NCLDVs. However, a few closely-related strains had drastically different seasonal dynamics, suggesting that while phylogenetic proximity often indicates ecological similarity, occasionally phenology can shift rapidly, possibly due to host-switching. Finally, we identified distinct functional content and possible host interactions of two major NCLDV orders-including connections of Imitervirales with primary producers like the diatom *Chaetoceros* and widespread marine grazers like *Paraphysomonas* and Spirotrichea ciliates. Together, our results reveal key insights on season-specific effect of phylogenetically distinct giant virus communities on marine protist metabolism, biogeochemical fluxes and carbon cycling.

## Introduction

Viruses are the most abundant and genetically diverse entities in marine systems [1, 2]. Orders of magnitude more abundant than prokaryotes and eukaryotes, viruses exert significant influence on ecosystem processes, such as structuring microbial communities and altering biogeochemical cycles [3, 4, 5]. Much of what is known about marine viruses stems from studying viruses that infect bacteria and a substantial knowledge gap exists regarding viruses that infect marine eukaryotes or how these eukaryotic-virus interactions modulate ecosystem processes. Globally, eukaryotes contribute significantly to ocean biomass and primary production [6, 7, 8], suggesting that eukaryotic viruses have the potential to exert significant control on marine systems.

Many of the viruses that infect marine eukaryotes belong to the phylum Nucleocytoviricota and are referred to as nucleocytoplasmic large DNA viruses (NCLDVs; [9, 10, 11]). This diverse monophyletic group is composed of several orders including Algavirales (Phycodnaviruses), Imitervirales (Mimiviruses), Pimascovirales, Pandoravirales, Chitovirales, and Asfuvirales, and infects many microbial eukaryotic lineages, including chlorophytes, haptophytes, dinoflagellates, choanoflagellates, and stramenopiles [12–18]. These viruses have large genomes up to 2.5 Mbp [19] harboring many genes for DNA replication, translation, and repair, as well as suites of genes implicated in the rewiring of host metabolic networks [20–22, 12]. Virus-encoded genes involved in nutrient acquisition and utilization, photosynthesis, and carbon fixation alter the growth of eukaryotic hosts and potentially also the nutrient cycles in which these hosts are embedded [16, 12, 21–27].

NCLDVs are widely distributed throughout the ocean and display distinct latitudinal patterns in community structure that correlate with eukaryotic communities [12, 28–31]. Yet, with a few notable exceptions like the extensive work on viruses infecting the coccolithophore *E. huxleyi* [32–38], our understanding of these viral communities as a whole is based on spatial surveys and short-term studies, with little existing data on the long-term temporal dynamics of these viruses; the latter is crucial to assess their ecological succession and impact on the microbial eukaryote populations. [39]. Nevertheless, we know that many microbial eukaryote populations exhibit strong temporal patterns (e.g., [40–43]), suggesting that viral communities should also demonstrate similarly variable patterns in incidence and abundance. Patterns of seasonal dynamics have already been elucidated in prokaryotic viruses [44] but a better understanding of the temporal dynamics of eukaryotic viruses and hosts is necessary to understand the impact of eukaryotic viruses on ecosystem processes.

We leveraged a multi-year, monthly metagenomic time series [44,45] to quantify the diversity of NCLDVs off the coast of Southern California and to characterize the temporal dynamics of these populations. Together, using this rich dataset we identified clusters of NCLDVs with distinct classes of seasonal and non-seasonal temporal dynamics and showed that closely related strains typically had similar dynamics. As an exception to this rule, in a select few cases, closely related strains had drastically different seasonal dynamics, suggesting that while phylogenetic proximity often indicates ecological similarity for marine NCLDVs, occasionally phenology can shift rapidly, possibly as a result of host-switching.

## Methods

### Sample Collection, DNA extraction, and sequencing

Surface seawater was collected monthly from 5 m as part of San Pedro Ocean Time-series (SPOT) (https://dornsife.usc.edu/spot/) using a 12 x12 Niskin bottle rosette directly into 20-L cubitainers. Seawater was filtered for viral metagenomes (< 0.22*µ*m, n = 53) by serially passing 0.5 to 1 L of seawater through an 80*µ*m mesh and a 0.2*µ*m Sterivex filter (Millipore) onto a 0.02*µ*m Anotop filter (Whatman). Similarly, DNA was collected for cellular metagenomes (0.22 - 1.2*µ*m, n = 42) and rRNA gene amplicons (1.2-80*µ*m, n = 46) monthly between May 2009 and August 2014 and May 2009 and June 2013, respectively, by passing 15 to 20 L of seawater through 80*µ*m mesh and a 1.2*µ*m AE filter and onto a 0.22*µ*m Sterivex filter.

Viral-fraction DNA was sent to JGI for library preparation and Illumina HiSeq-2000 2 x 150 bp sequencing under CSP proposal 2799 [44–45]. DNA for cellular metagenomes and cellular 18S rRNA gene amplicons were extracted using the Qiagen AllPrep DNA/RNA kit (Qiagen) and the DNA was cleaned and concentrated using AMPure beads (Beckman Coulter). 18S rRNA amplicons were amplified using universal primer sets 515Y and 926R as described by Yeh et al [46]. Cellular fraction metagenomic libraries were prepared and sequenced as described previously in Dart et. al. [45].

### Read quality control and assembly

Viral and cellular metagenomic assemblies for each sample were downloaded from the JGI genome portal (proposal 2799). Reads from both metagenomes were subject to trimming using bbtools [47], read correction using bfc [48], and assembly using SPAdes [49] with a range of k-mers and the following options: ‘-m 2000 --only-assembler -k 33,55,77,99,127 --meta -t 32’.

18S rRNA amplicon sequences were trimmed using cutadapt [50] and amplicons were pooled using bbsplit with the PR2 database [51]. The 18S rRNA amplicons were analyzed using DADA2 [52] implemented in QIIME2 [53]. 18S rRNA reads were then denoised and filtered for chimeras using DADA2 commands and assigned taxonomy against the PR2 database [51].

### Identification of NCLDV phylotypes

To identify NCLDV phylotypes, protein-coding regions for assemblies were predicted using Prodigal [54], and the protein sequences were searched for copies of the DNA Polymerase Beta (PolB) gene using hmmsearch (e-value 10*^−^*^5^) in HMMER3 [55] using the polB model from Moniruzzaman et al. [26]. PolB sequences were dereplicated at 99% amino acid identity using cd-hit [56,57] requiring 90% coverage. Only sequences greater than 500 amino acids in length (seqkit; [58]) were considered for downstream analyses. PolB sequences were classified using GraftM [59] to remove any non-NCLDV PolB sequences.

Reads were mapped to the corresponding PolB nucleotide sequence using BWA-MEM [60] and sorted and indexed with samtools [61, 62]. Read coverage was calculated using ‘trimmed_mean’ in CoverM (v0.7) (https://github.com/wwood/CoverM) requiring 98% minimum read identity.

### Metagenomic binning, quality control, and annotation

Metagenomic reads were aligned to contigs using BWA-MEM and sorted and indexed with samtools [61, 62]. Bins were assembled using MetaBat 2 [63] with ‘runMetaBat.sh -s 100000 -m 5000 –minS 75 –maxEdges 75’. NCLDV bins were identified and scanned for completeness using miComplete [64] using five conserved NCLDV marker genes, namely A32 ATPase, Major Capsid Protein (MCP), SFII helicase, B-family DNA polymerase (PolB), and VLTF3 transcription factor [26]. NCLDV bins with ≥ 3 of the five marker genes were retained for downstream analyses. NCLDV bins were screened with Viralrecall [65] using the NCLDV-specific database (NVOG), requiring a minimum of 3 hits to the NVOG database, and a minimum score of 0. Bins were dereplicated with dRep [66] at 95% nucleotide identity requiring coverage of 30% the shorter bin. Protein coding regions for the NCLDV bins were predicted using Prodigal [54] and annotated using eggNOG-mapper [67,68] using default settings.

VirHostMatcher [69] was used to predict putative hosts of the NCLDVs based on oligonucleotide frequency patterns. Reference transcriptomes from the Marine Microbial Eukaryote Transcriptome Sequencing Project (MMETSP; [70, 71]) were used as the host database and d2* dissimilarity was calculated using the NCLDV bins. Here we report the lowest d2* dissimilarity for each bin below 0.4. To supplement this approach, a co-occurrence network was created to link NCLDV phylotypes to eukaryotic OTUs using FlashWeave [72]. The resulting correlations were filtered using a weight cutoff of 0.4 [73] and displayed using Cytoscape [74].

### Phylogenetic analyses of NCLDV marker genes

PolB phylotype sequences were aligned using mafft [75, 76] with ‘–retree 2’ and alignments were trimmed using trimal [77] with the ‘-gappyout’ flag. A maximum likelihood tree of PolB phylotypes was constructed using RAxML [78] with ‘raxmlHPC-PTHREADS-SSE3 -k -T 80 -f a -m PROTGAMMAGTR -p 12345 -x 12345’.

A multigene tree was constructed using a concatenated alignment of five conserved NCLDV marker genes (A32, MCP, SFII, PolB, and VLTF3) using ncldv_markersearch.py [26]. Poorly aligned sequences and regions were removed using trimal [77] with ‘-gappyout’. A maximum likelihood tree was constructed using RAxML [78], specifically ‘raxmlHPC-PTHREADS-SSE3 -k -T 80 -f a -m PROTGAMMAGTR -p 12345 -x 12345’. This tree was rooted using Chitovirales as an outgroup.

### Time-series analysis and clustering

Bray-Curtis similarity was calculated using the *distance* function in the phyloseq R package (v3.19), [79]). Seasonal profiles were calculated as trimmed mean coverage normalized by the number of reads in each metagenomic sample for NCLDV phylotypes across samples. For host 16S rRNA and 18S rRNA sequences, relative abundance was used. Finally, data was linearly interpolated to create an evenly spaced time series with no missing data using the *ApproxFun* in the R stats package. Spectral densities were calculated using *spectrum* in the stats R package.

Using the time series generated above, *ptestg* in the ptest package [80, 81] using the method “extendedRobust” was applied to identify periodicity in these time series with a p-value cutoff of *p <* 0.001. Time series were clustered using z-score normalized monthly mean seasonal profiles with the function *repr_seas_profile* in the TSrepr package [82, 83]. The number of clusters was determined based on the minimum Davies-Bouldin index value [84].

Visualization was performed using a combination of functions from the ggplot2 [85], ggtree [86], and ggpubr [87] R packages as well as base R [88].

### Data availability

The viral metagenomes are available at the JGI Genome portal under proposal ID 2799. Cellular metagenomes are available at the National Center for Biotechnology Information (NCBI) under BioProject PRJNA814250 (SRR18278848-SRR18278886) and 18S rRNA amplicons are available on EMBL-EBI under projects PRJEB48162 and PRJEB35673. NCLDV MAGs, PolB phylotypes, protein annotations, host prediction data, and phylogenetic trees are available in a Figshare repository (https://figshare.com/projects/Seasonal_Dynamics_of_Giant_viruses_in_Long-Term_metagenomic_time_series_SPOT_/206872).

## Results

### High NCLDV Prevalence and Diversity in the California Current

We found a diverse set of NCLDV phylotypes on the basis of the reconstructed PolB marker gene. We use the term phylotype throughout the text to indicate reconstructed NCDLV PolB gene sequences clustered at 99% amino acid identity. A total of 676 NCLDV phylotypes were identified in the two size fractions: 655 phylotypes were found in both the viral and cellular size fractions, and 15 and 6 phylotypes were found exclusively in the viral and cellular size fractions, respectively. The NCLDV community was dominated by Algavirales (59%) and Imitervirales (36%), with Pimascovirales having a minor contribution (0.7%) (Figure 1). This is consistent with a recent global survey that found Imitervirales and Algavirales to be the two dominant NCLDV orders in the world’s oceans [12]. Imitervirales were more abundant in the larger size fraction (0.2 - 1.2*µ*m) compared to the viral size fraction (0.02 - 0.2*µ*m), consistent with the often-large size of their viral particles [89].

**Figure 1.**
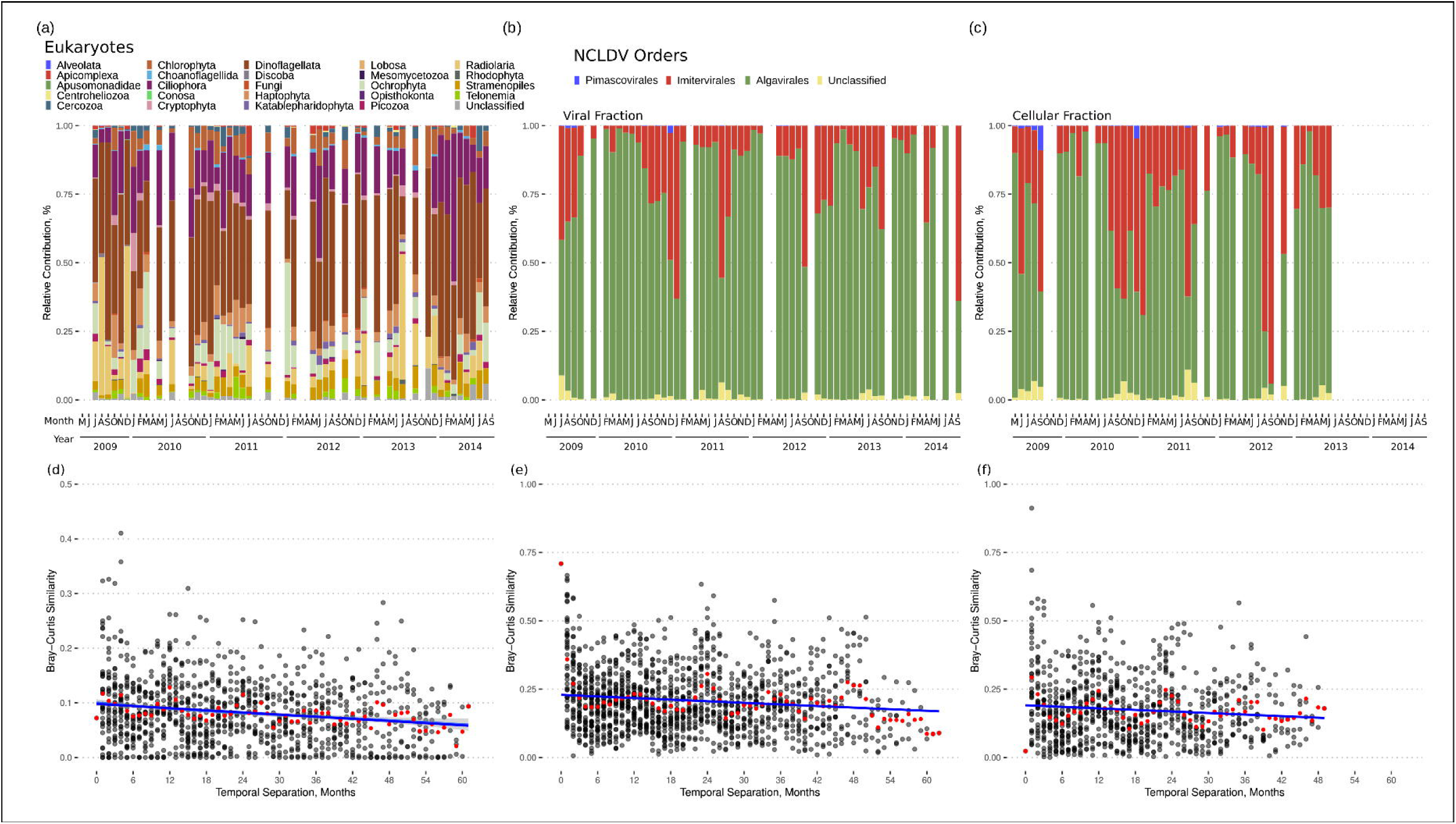
NCLDV and host communities at SPOT vary seasonally in their composition. **(a,d)** The eukaryotic community assayed via 18S rRNA amplicon sequencing in terms of (a) the overall composition and (d) pairwise community similarity calculated between all sample pairs. (b,e) The relative abundance of NCLDV orders in the viral size fraction (0.02-0.2μm) was assayed via reconstructed PolB sequences. (c,f) The same was done for the cellular size fraction (0.2-1.2μm). In panels (d-f) red lines indicate the mean Bray-Curtis similarity for a given temporal separation.

The eukaryotic community, based on 18S rRNA genes in the eukaryotic size fraction (1.2 - 80 *µ*m), was dominated by Dinoflagellata and Ciliophora, with 18S rRNA capturing both heterotrophic and phototrophic eukaryotes. Eukaryotes in the smaller cellular fraction (0.2 - 1.2*µ*m) were detected using chloroplast 16S rRNA. Eukaryotes in the Mamiellaceae order were abundant in the smaller cellular size fraction, including *Bathycoccus, Micromonas,* and *Ostreococcus*. Bacillariophyta and Prymnesiophyceae were also abundant in the eukaryotic size fraction (Figure S1).

### NCLDV Populations Frequently Exhibit Strong Seasonal Dynamics

We quantified the relative coverage of 676 NCLDV PolB phylotypes across our five-year metagenomic times series in two separate filtered size fractions (0.02-0.2*µ*m “viral” fraction and a 0.2-1.2*µ*m “cellular” fraction). Overall, the NCLDV community exhibited seasonally predictable changes in composition (Figure 1), with peak community similarity (Bray-Curtis) occurring at 1-year intervals, in both sequenced size fractions. Cycles were apparent at the order level with Imitervirales dominating in the late summer and early fall and Algavirales dominating at all other times. These cycles were apparent, though less clearly so, in the corresponding eukaryotic community (Figure 1a, d). Among the NCLDV, the community found in the larger size fraction (0.2-1.2*µ*m, “cellular”; Figure 1c, f) had stronger apparent fluctuations in composition than that found in the smaller fraction (0.02-0.2*µ*m, “viral”; Figure 1b, e). This difference is likely due to the fact that many of the free-floating NCLDVs have sufficiently large viral particles that they are likely to not be found in the smaller “viral” fraction. NCLDV particles at SPOT might be present in higher proportion in the cellular size fractions, contributing to the relatively stronger annual periodicity observed within this fraction. [19, 89–91].

While overall community similarity showed seasonal fluctuations, it was initially unclear if these were driven by fluctuations in the abundance of all or a subset of community members. We individually examined the temporal patterns in abundance of each NCLDV to classify strains as seasonal or non-seasonal (Figure 2), and to distinguish strains with annual peaks from those that peaked biannually. Using a likelihood-ratio test to compare fitted harmonic models we found 13 NCLDVs with strong seasonal dynamics. The same strains were recovered as significantly seasonal using Fisher’s g test. We then applied a robust variant of Fisher’s g test that recovered a greatly expanded set of significantly periodic strains (131). Many of these strains had unevenly sized annual peaks or “gap” years in which they did not peak, possibly explaining why less robust methods failed to recover them. Even among the strains in which no seasonality was detected using the more robust test, it was clear that some groups of strains showed some apparent seasonal dynamics, in that their peaks always occurred during a specific season even if they did not peak every year (Figure S2). This suggests that longer time series studies may have the power to detect the many potentially seasonal viral populations with a patchy distribution across years. Overall, viruses could be classified as (1) seasonal with one annual peak (17%), (2) seasonal with two annual peaks (corresponding to spring and fall upwellings) (2%), or (3) erratic with one or a couple of peaks seemingly randomly distributed across years (81%).

**Figure 2:**
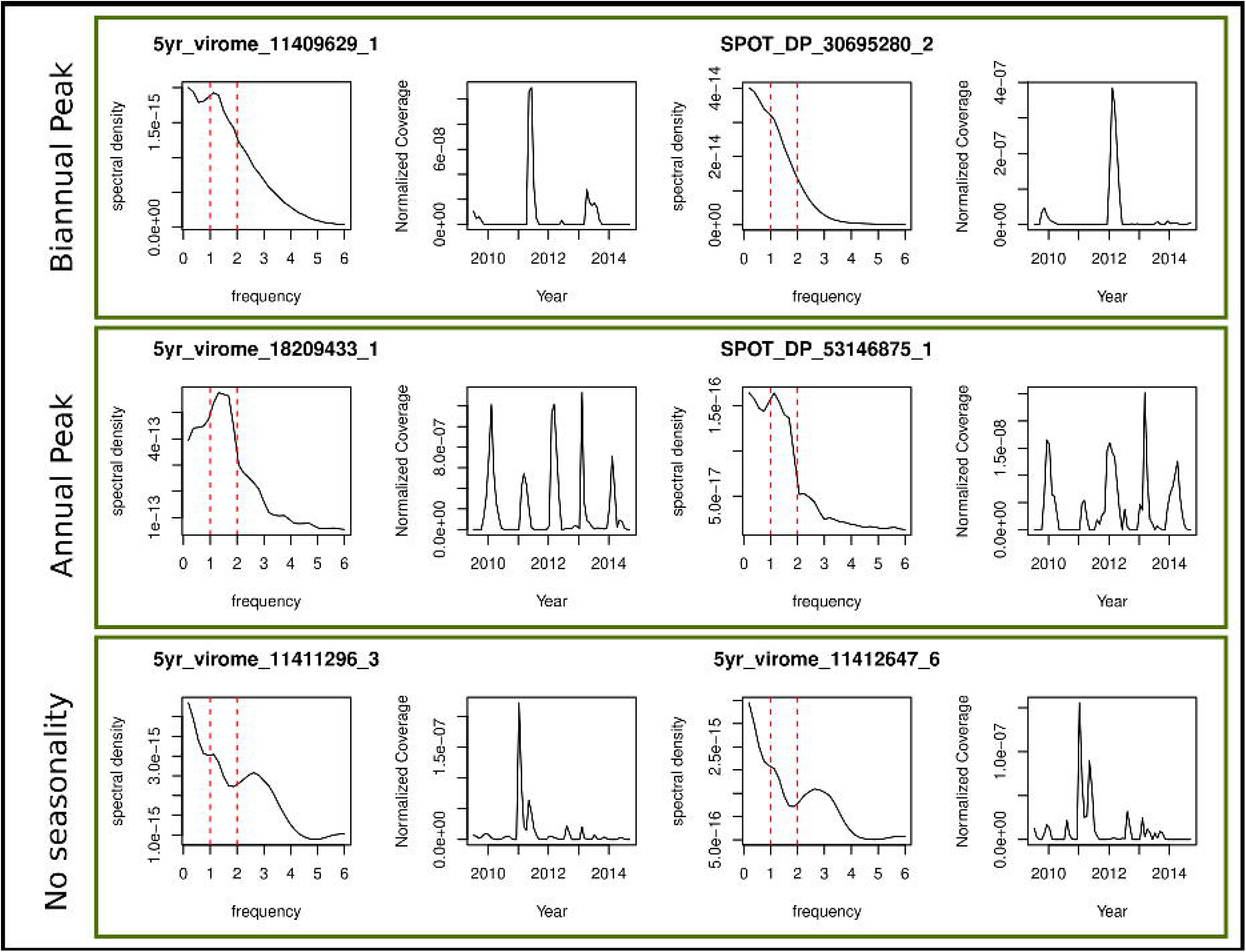
Periodograms to detect seasonality in NCLDV time series. Periodograms represent a decomposition of the time series into a set of periodic functions whose frequencies are represented in the periodogram. High spectral density indicates the dominant frequencies in a time series, expressed in units of 1/years. The class of statistical tests applied in the main text calculates various summary statistics based on these periodograms to assess the significance of seasonal patterns. Here we show representative periodograms along with corresponding normalized coverage for viruses with biannual peaks, annual peaks, and no seasonal peaks.

### Closely Related NCLDVs Have Similar Seasonal Dynamics

We clustered phylotypes that exhibited a single annual peak into distinct seasonal clusters based on their mean seasonal profile (Figure 3). NCLDV phylotypes were clustered in each size fraction separately resulting in 10 and 9 clusters in the viral and cellular size fraction, respectively (Figure 3a, b). The clusters showed a succession of NCLDV phylotypes throughout the year, with 11 NCLDV peaking in spring (Mar-May), 7 peaking in summer (Jun-Aug), 21 peaking in fall (Sept-Nov), and 31 peaking in winter (Dec-Feb).

**Figure 3.**
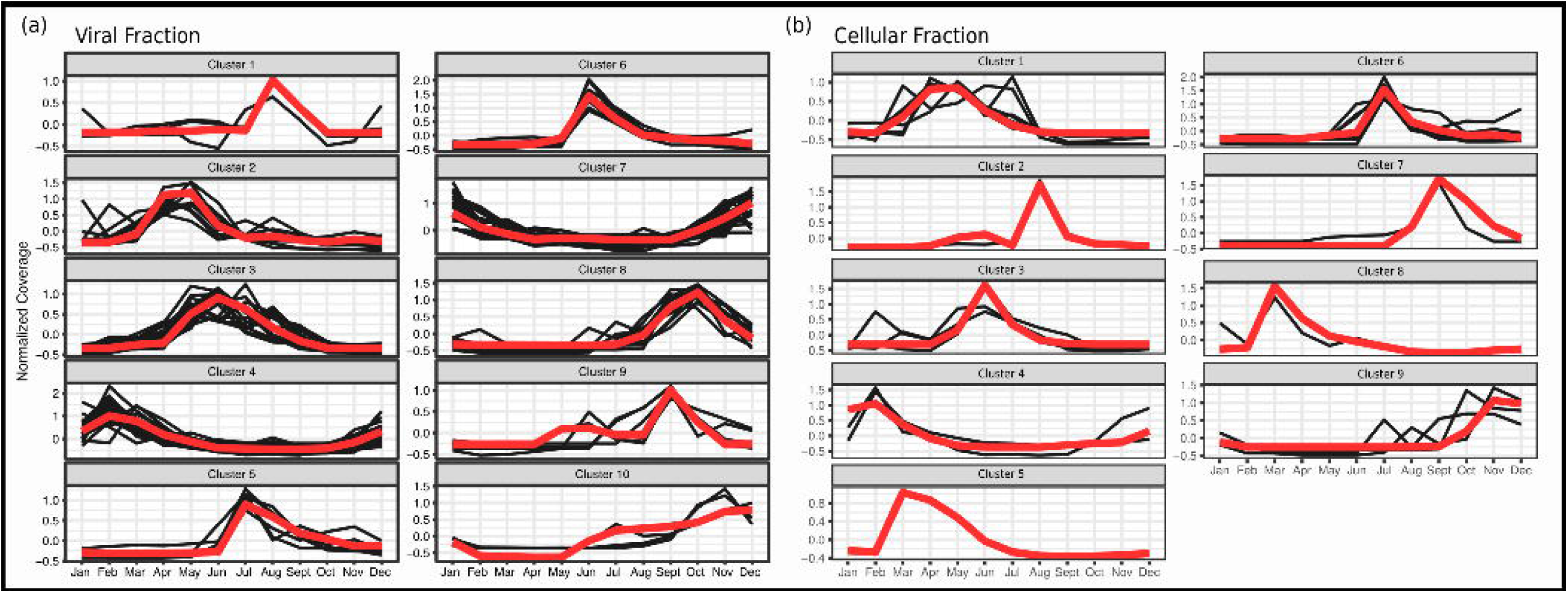
NCLDV phylotypes fall into distinct seasonal clusters. Distinct clusters for the (a) viral and (b) cellular fractions were calculated for significantly seasonal NCLDV phylotypes using k-medoids clustering and the minimum Davies-Boudin index. Dark lines indicate normalized mean coverage of an individual phylotype in the cluster and the medoid of each cluster is shown in red.

Phylogenetically related phylotypes tended to belong to the same or similar seasonal cluster (Figures 4-5). For pairs of phylotypes, the phylogenetic distance was negatively related to the agreement of the pair’s seasonal abundance profiles (Figures 4c,5c). Prasinoviruses (Algavirales) typically belonged to the spring and early summer clusters, while Imitervirales belonged to clusters peaking in later summer and fall. Prasinoviruses were also hyperabundant compared to other NCLDV phylotypes (Figure 4b,5b). Interestingly, Algavirales typically peaked earlier (February) in the viral size fraction and later (April) in the cellular size fraction (Figures 4a-5a), suggesting that larger viruses peak later into the spring in response to their algal hosts blooming during this season [92,93].

**Figure 4.**
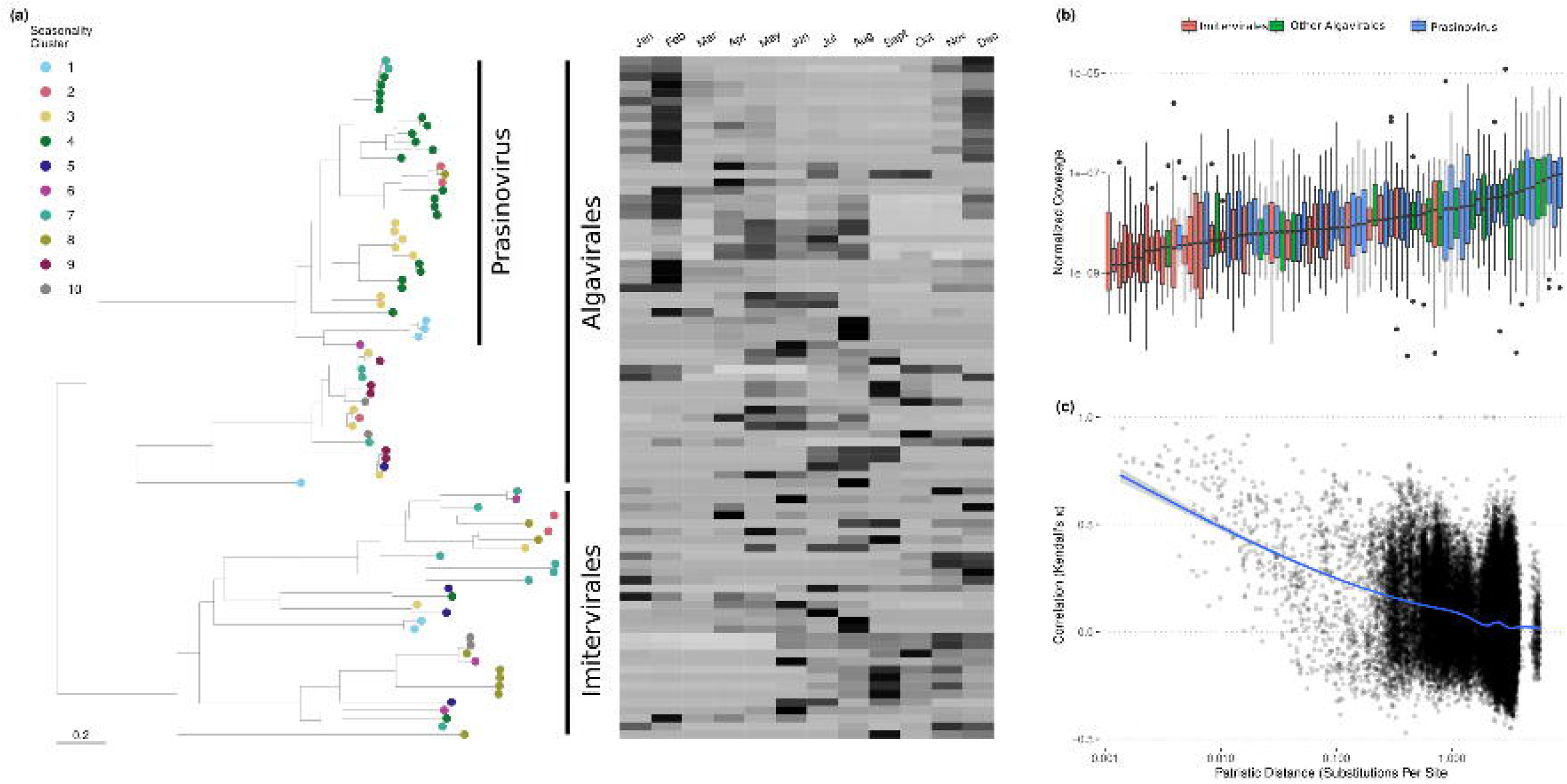
Closely related phylotypes tend to have similar seasonal dynamics (viral size fraction; 0.02-0.2μm). (a) Phylogeny of NCLDV PolB genes and mean annual abundance profiles for each gene. (b) Abundance of each phylotype across samples. Note that Prasinoviruses are particularly abundant in this system. (c) Pairwise correlations in phylotype abundances across time plotted against patristic distances computed from the phylogeny in (a). An expanded figure with phylogenetic tree leaf labels is available in Figure S3.

**Figure 5.**
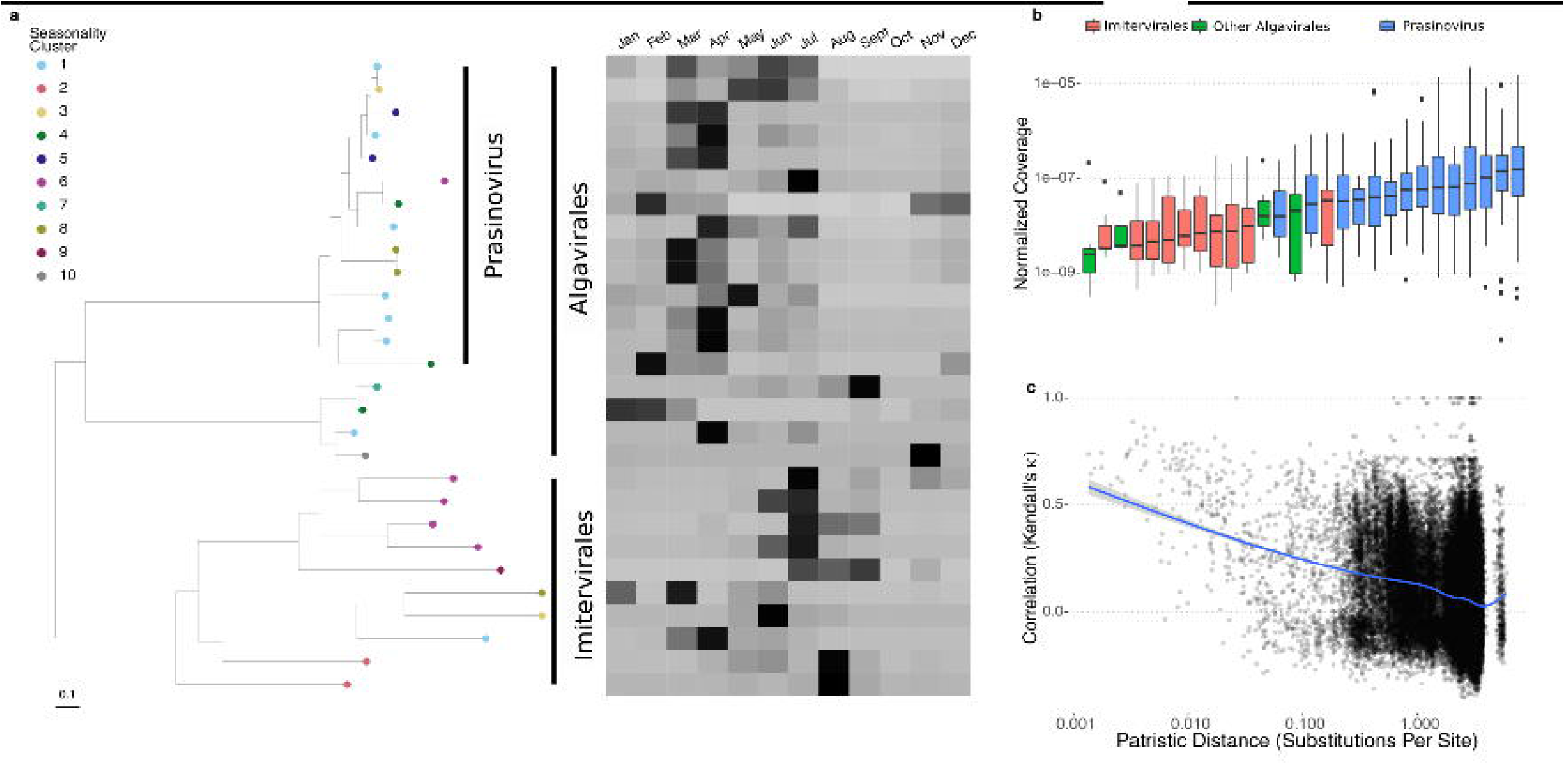
Closely related phylotypes tend to have similar seasonal dynamics (cellular size fraction; 0.2-1.2μm). (a) Phylogeny of NCLDV PolB genes and mean annual abundance profiles for each gene. Observe that peak abundances for Prasinoviruses are shifted later in the year relative to those shown in the viral fraction in Fig 4. (b) The abundance of each phylotype across samples. (c) Pairwise correlations in phylotype abundances across time plotted against patristic distances computed from the phylogeny in (a). An expanded figure with phylogenetic tree leaf labels is available in Figure S4.

Finally, we note that even though closely related strains tend to have similar seasonal dynamics, occasionally a pair of closely related strains would have drastically different seasonal profiles (Figures 4a-5a). These changes could indicate host shift events and suggest that high-resolution time-series data may be a useful tool to detect viral host shifts along a phylogeny. Alternatively, it is also possible that some protist lineages within a phylogenetically cohesive group showed divergent seasonal dynamics. If the viruses associated with these protists tracked their dynamics, that could at least partly explain our observations.

### Eukaryotic Hosts Exhibit a Weaker Seasonality than Their Viruses

We found far fewer eukaryotic taxa had detectable seasonal dynamics than we found for NCLDVs (18) consistent with overall weaker seasonal fluctuations found at the community level (Figure 1). While initially surprising, we suspect that the relatively low population densities of the hosts relative to the prokaryotic communities sequenced simultaneously may mean noise masks potential seasonal patterns. Ultimately, eukaryotic taxa clustered into fewer (3) seasonal clusters, mainly corresponding to the major annual upwelling events: early spring, late spring, and fall (Figure S5).

### Distinct Functional Potentials Among Two NCLDV Families

Adding metagenomic bins to the phylotype analyses provided additional information regarding the NCLDV community, including genomic potential and putative hosts. Seventy NCLDV metagenome assembled genomes (MAGs) were assembled with ≥ 3 marker genes and were used for subsequent analyses. The MAGs ranged in size from 72-826 kb and contained 100-857 coding sequences, MAG GC content ranged between 20.2 and 54.3% (Figure S6). A multigene phylogenetic tree indicated that 31 MAGs belong to Algavirales, 38 MAGs to Imitervirales and one was Pimascovirales (Figure 6a). According to eggNOG annotations, Imitervirales MAGs had a particularly large number of genes belonging to COG functional groups, including DNA replication, translation, transcription, and cell wall envelope biogenesis (Fig 6b).

**Figure 6:**
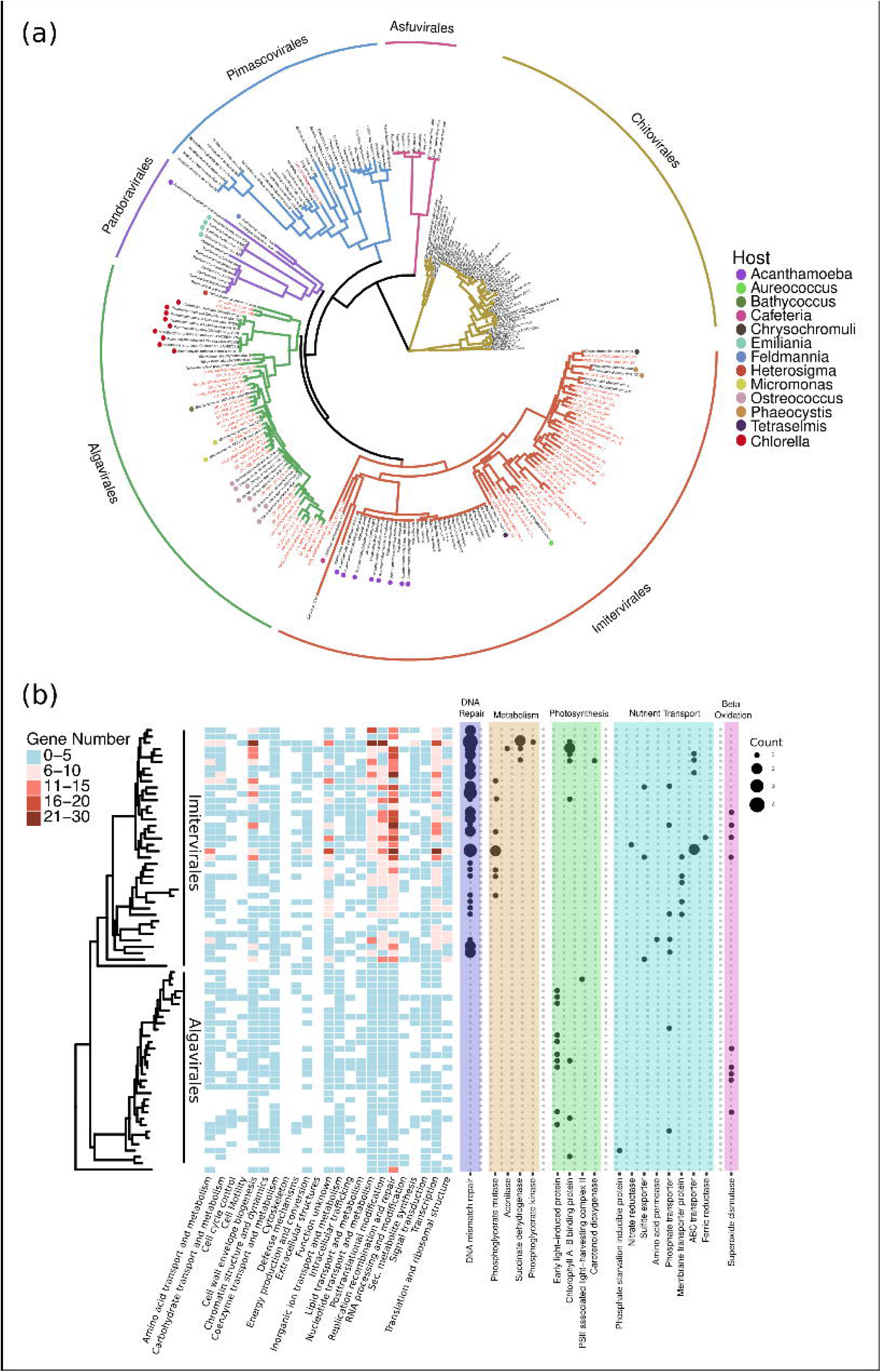
Phylogenetic and functional potential of NCLDV MAGs. (a) Multigene Maximum likelihood phylogenetic tree from a concatenated alignment of five conserved NCLDV genes from reference genomes (black) and MAGs (red). The node color indicates the reported host of reference NCLDVs, and the branch color indicates NCLDV order membership. (b) The tree from (a) subset to include only MAGs generated here. The heatmap (center) indicates the number of genes within each COG functional group and the bubble plot (right) indicates the number of genes involved in select metabolic functions.

NCLDV MAGs also encoded diverse auxiliary metabolic genes (AMGs) homologous to sets of AMGs previously identified among viral genomes. There were clear order-level differences in the sets of AMGs encoded between Imitervirales and Algavirales MAGs (Figure 6b).

Imitervirales MAGs encoded a diverse set of genes involved in energy metabolism and transport, which were all but absent among Algavirales MAGs. This is consistent with previous work showing that Imitervirale members encode diverse transporters both for the transport of dNTPs across the mitochondrial membrane [94] and for the transport of nutrients across the cell membrane [95]. Surprisingly few Algavirales MAGs from our study site encoded nutrient transporters, despite the apparent importance of Algavirales-encoded transporters in some other systems [95, 21]. This may suggest less nutrient limitation of Algavirus replication at SPOT compared to other previously studied locations, which are often more oligotrophic; and we note Algaviruses are less abundant at SPOT in the more oligotrophic late summer and fall seasons. While both Imitervirales and Algavirales MAGs encoded light-harvesting orthologous groups of genes, each NCLDV order encoded distinct sets with specific functions, suggesting that these two orders modulate host photosynthesis using distinct pathways. Algavirales MAGs encoded many orthologous groups directly implicated in photosynthesis, whereas such groups were nearly absent among Imitervirales MAGs. On the other hand, several Imitervirales MAGs, as well as several Algavirales MAGs, encoded various chlorophyll-binding proteins. Finally, superoxide dismutases were found distributed across both Imitervirales and Algavirales MAGs, suggesting that superoxides produced by host metabolism or stress responses may pose a general threat to infecting NCLDVs.

### Compositional Patterns Reveal Putative NCLDV Hosts

We identified putative host pairings for our NCLDV MAGs based on k-mer frequency and phylotypes based on a co-occurrence network (Fig 7a, b). Both methodologies allowed for the prediction of a diverse set of hosts for our recovered NCLDVs although we acknowledge that correlation and sequence-based methods have limitations and may sometimes yield false positives, however, they remain valuable tools for identifying valid host-virus pairings.

**Figure 7:**
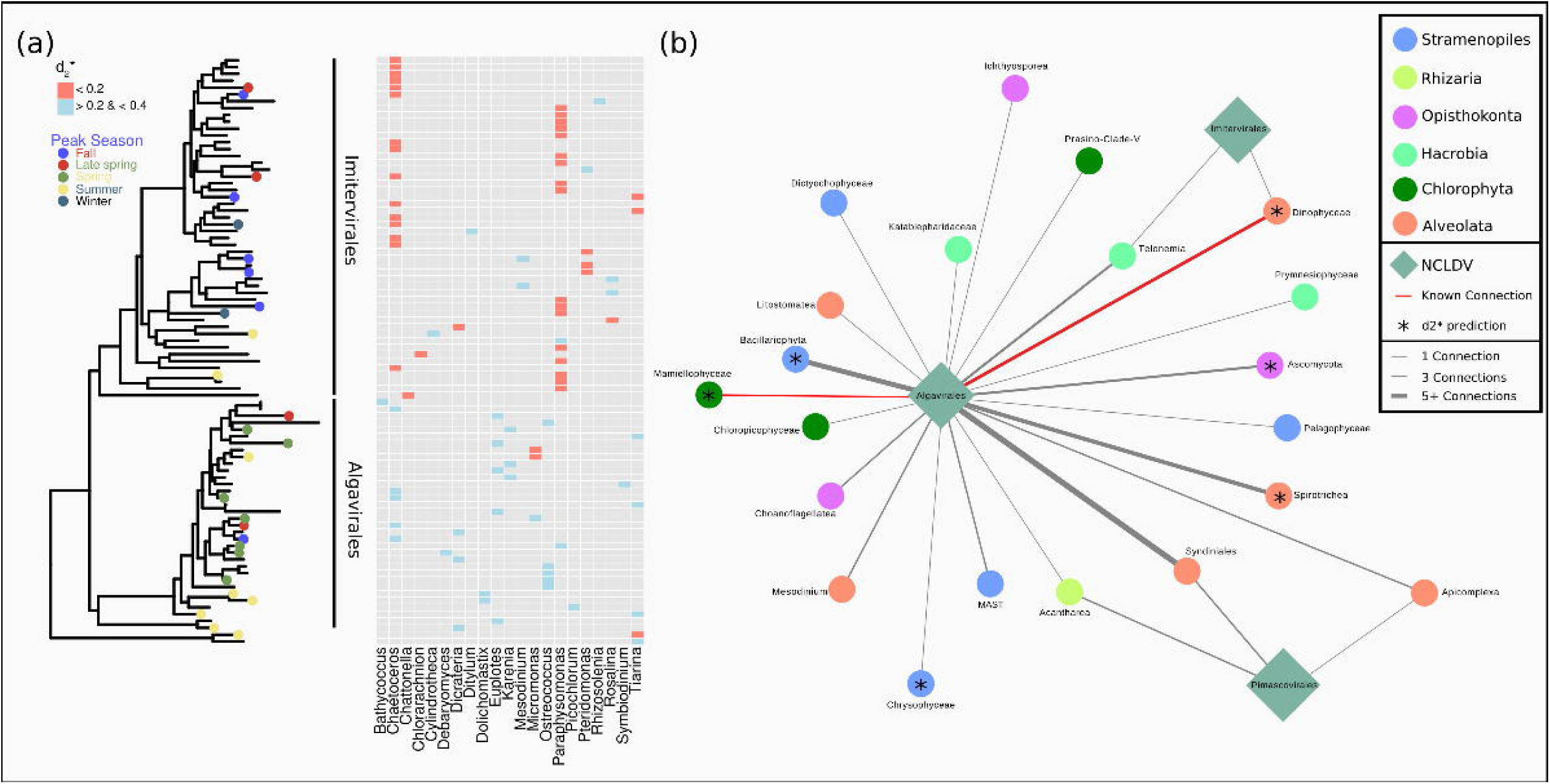
Reconstructed NCLDV MAGs are predicted to infect diverse host taxa. (a) Phylogenetic tree of NCLDV MAGs and their predicted host taxa based on k-mer profile similarity (d2∗) with MMETSP reference genomes. High-confidence (d2∗ < 0.2) host predictions are shown in red, while lower-confidence host predictions are shown in blue. Peak season is also shown for MAGs with significant seasonality. (b) Co-occurrence network of NCLDV PolB phylotypes and eukaryote OTUs using 18S data. A weight cutoff of 0.4 was used to filter connections, and the number of connections is shown by the edge thickness. Known NCLDV and eukaryotic host pairs from cultured isolates are shown in red and connections match and d2* results are shown with a star (*). An expanded network and phylogenetic tree are available in Figure S7.

Correlation-based methods have been used previously to successfully recover established giant virus-host linkages in different marine environments [96]. We report many putative connections between Imitervirales MAGs, which were more abundant during the warmer and more oligotrophic late summer and fall seasons, and *Chaetoceros* and *Paraphysomonas* genera (Figure 7a). *Chaetoceros* is a diverse genus of marine diatoms and is globally distributed [97, 98], yet this is the first indication, to our knowledge, of an NCLDV infecting a member of this genus.

*Paraphysomonas* is a cosmopolitan phagotrophic clade [99] belonging to the chrysophytes, which were recently suggested to be a target of many novel Imitervirales lineages based on host-virus correlations in data from the Tara Oceans Project [12]. Relatively few hosts for Algavirales MAGs could be predicted with high confidence, but those that could belong to the genus *Micromonas,* a common, sometimes hyperabundant [100] marine photosynthetic picoeukaryote, occurring year-round at SPOT.

Additional to the k-mer frequency-based method, the co-occurrence approach allowed for the prediction of environmentally specific hosts for all three orders of recovered phylotypes (Figure 7b). Many of these connections (e.g., Mamiellophyceae: *Micromonas*) were consistent with results from the k-mer frequency approach, adding confidence to these predictions. A large number of associations were found between Algavirales and Syndiniales. In addition, we also recovered associations between Algavirales members and Dinophyceae, consistent with previous observations [101]. Finally, connections were also observed between Algavirales and Spirotrichea, which are widespread ciliates and a major trophic link from nanoplankton to higher trophic levels [102].

## Discussion

We leveraged a high-resolution multi-year metagenomic time series to assess seasonal patterns of NCLDV abundance in the California Current. This work demonstrates the advantages of long-term sequencing projects that sample multiple size fractions and have frequent sampling [44–46]. We found strong seasonal patterns in NCLDV phylotype abundance that differed systematically across size fractions, consistent with patterns seen for phage infecting cyanobacteria (cyanophage) in these same sample sets [45]. By incorporating multiple years of data in our analysis we were able to detect significant seasonal patterns even in the presence of abnormal years where a peak in abundance did not occur or was shifted.

While previous work has demonstrated the global distribution and numerical abundance of marine NCLDVs [12], these explorations have only captured a snapshot of NCLDV dynamics. By leveraging multi-year data, we show that these viral populations are extremely dynamic in time, often varying to a greater degree than even their putative hosts (Figures 1-5). The majority of phylotypes exhibit no seasonality and are apparently erratic in their patterns, similar to what has been seen for species-like populations of cyanophage at SPOT [45]. Phylotypes that are seasonal peak at different times, presumably following when their particular hosts bloom in abundance, a pattern also paralleled in cyanobacteria-cyanophage dynamics here [45].

Biogeographic investigations must consider this time-varying quality of NCLDV populations moving forward, as it is likely that many local populations vary in time. At the same time, this variation is non-random, as the community returns to a similar compositional state at one-year increments (Figure 1). The strong seasonality in the overall community of NCLDV follows similar patterns documented for phage, prokaryotic, and eukaryotic components of the microbial communities at SPOT [45–46, 103–105], emphasizing the interconnectedness of these communities. As global climate shifts, continued metagenomic surveillance will allow us to assess how seasonal dynamics change in marine microbial communities as environmental conditions are altered [106].

We found that NCLDV phylotypes group into distinct seasonal clusters, and that closely related phylotypes tend to share a cluster (Figures 4-5). This suggests that seasonal clusters reflect overall patterns in host identity since closely related strains are expected to share host taxa (Figure 7). Intriguingly, these patterns mean that temporal patterns of viral abundance may be used to place host-shift events on a viral phylogeny, independent of any knowledge about the host identity of any particular viral strain. Analysis of cyanophage similarly found that while phylogenetic proximity often indicates ecological similarity, closely related taxa sometimes differ in their phenology, indicative of host-switching. Thus, only small changes in genome sequences likely can drive shifts in the host range [107].

Finally, we found that the two major NCLDV orders found at our study site, Imitervirales and Algavirales, had distinct functional potentials based on an analysis of reconstructed viral MAGs (Figure 6). These two orders have distinct seasonal abundance patterns (Figure 1), suggesting that the functional composition of the NCLDV community fluctuates over time and phylogeny and functional potential are key drivers of NCLDV dynamics. These patterns may ultimately be caused by different ecological strategies between these two orders of NCLDV, but further research is needed to validate these patterns. It is not yet clear how different AMGs may play a role in the seasonal variation of viral communities [12, 22, 26], but we believe this will be a fruitful area of research moving forward.

In closing, this time-series analysis of NCLDV not only yields insight into their temporal dynamics but also genomic differences across major orders of these viruses. The distinct phenologies of NCLDVs likely reflect how they ‘track’ with the particular hosts they infect, and we leverage this to identify putative new host connections, including a novel possible connection to the diatom phylum Bacillariophyta. We emphasize that given dynamic changes in viruses over time biogeographic studies need to account for this in their interpretation of patterns. As sequencing costs continue to decline [108], and new tools for automatic sampling are developed [109], we expect the field to increasingly move towards building a high-resolution understanding of viral populations and evolutionary dynamics in the environment.

## Acknowledgments

This research was funded by the Joint Genome Institute (JGI), Simons Collaboration on Computational Biogeochemical Modeling of Marine Ecosystems / CBIOMES) grant [549943 to J.A.F.], Simons Award #653212 (JLW), Simons Award, NSF grants [EF-2125142 to J.A.F. NSF-OCE-1737409 to J.A.F.]; National Institutes of Health (NIH) grant 1R01GM120624-01A1 to N.A.A. and J.A.F. NSFC grant [42276163 to S.H.]; Shenzhen Science, Technology and Innovation Commission Programme [JCYJ20220530115401003 to S.H.]. M.M. and B.M. are supported through intramural funds provided by the Rosenstiel School of Marine, Atmospheric, and Earth Sciences, University of Miami.

## Conflict of Interest Statement

The authors declare no conflict of interest.

## Author contributions

JF, SH and MM supervised the research. JF secured external funding. SH designed and developed the NGS data analysis workflow. SL, JLW, SH and BM performed data analysis and writing of the manuscript. NA coordinated collaboration and data management and edited the manuscript. All authors critically reviewed and edited the manuscript.

## Notes

https://figshare.com/projects/Seasonal_Dynamics_of_Giant_viruses_in_Long-Term_metagenomic_time_series_SPOT_/206872

### Competing Interest Statement

The authors have declared no competing interest.

