## Supplementary information for "Phylogenetic proximity drives temporal succession of marine giant viruses in a five-year metagenomic time-series"

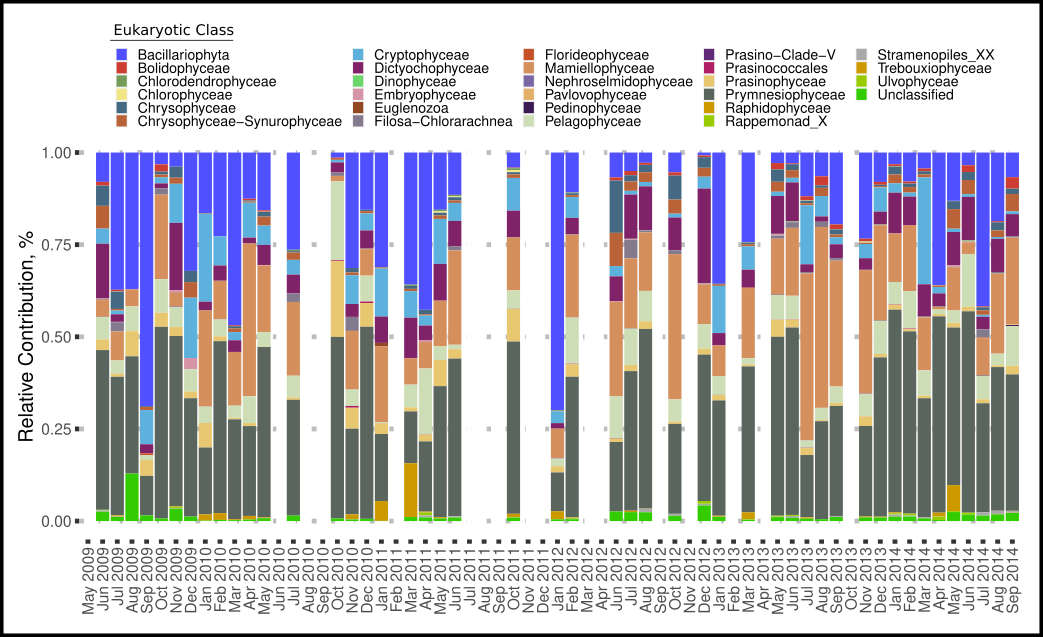

**Figure S1.** Eukaryotic diversity recovered from chloroplast 16S. The relative abundance of Eukaryotic OTUs classified to the class level is shown here.

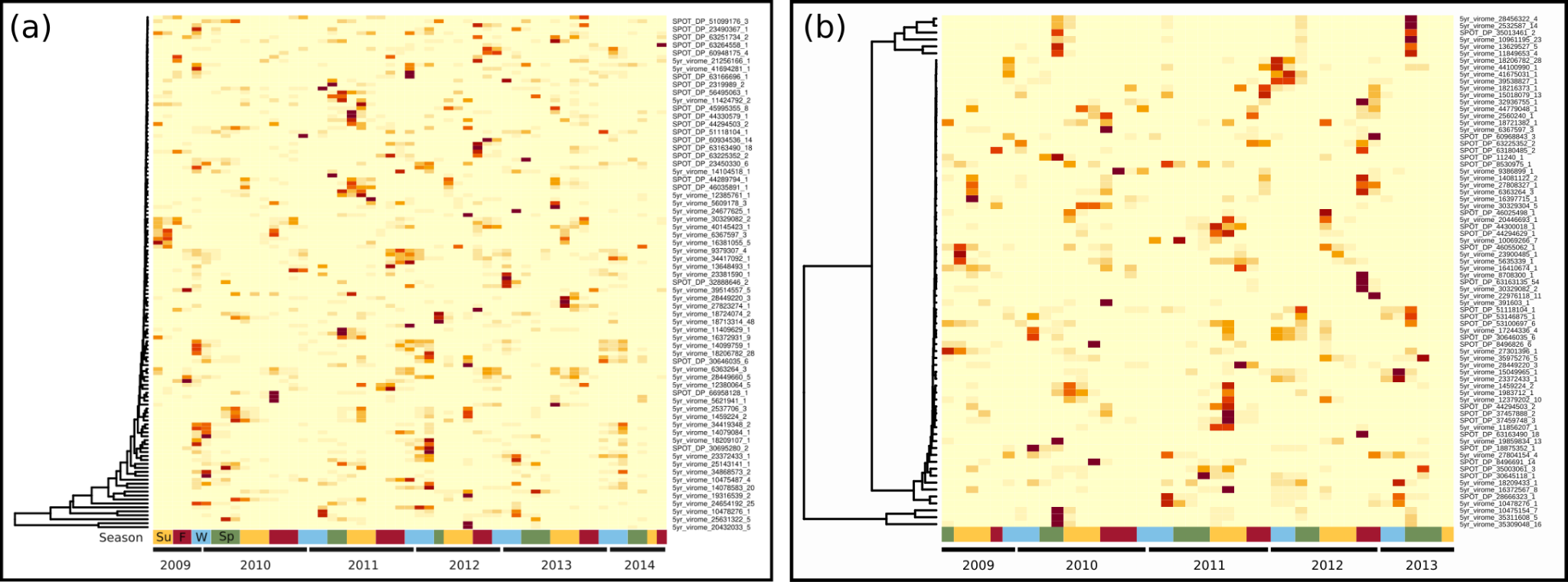

**Figure S2. Seasonality of NCLDV phylotypes.** Normalized abundance of the NCLDV PolB phylotypes with significant seasonality using the extended Fisher’s g test is shown in both the (a) viral and (b) cellular fraction.

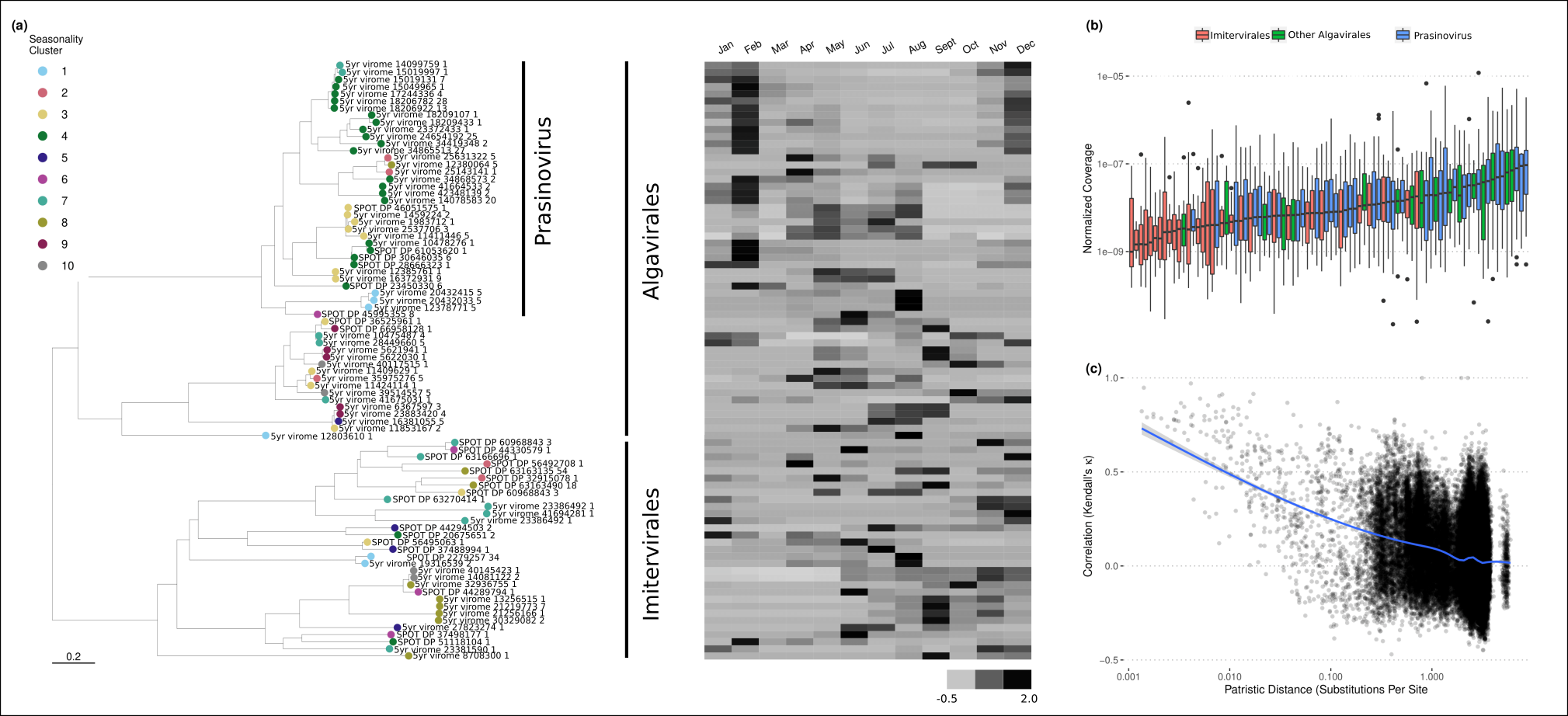

**Figure S3. Expanded figure 4.** Replica of figure 4 with leaf labels added to the phylogenetic tree.

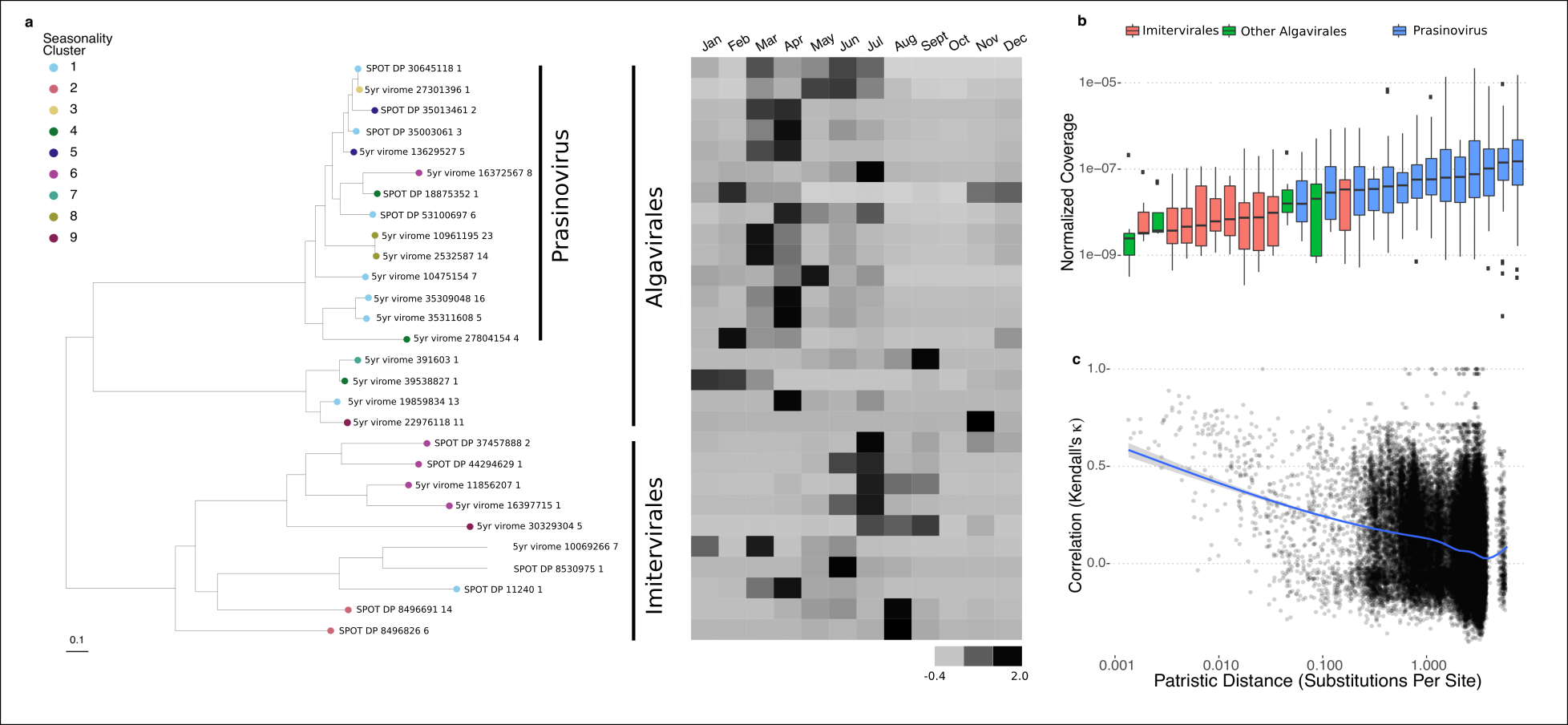

**Figure S4. Expanded figure 5.** Replica of figure 5 with leaf labels added to the phylogenetic tree.

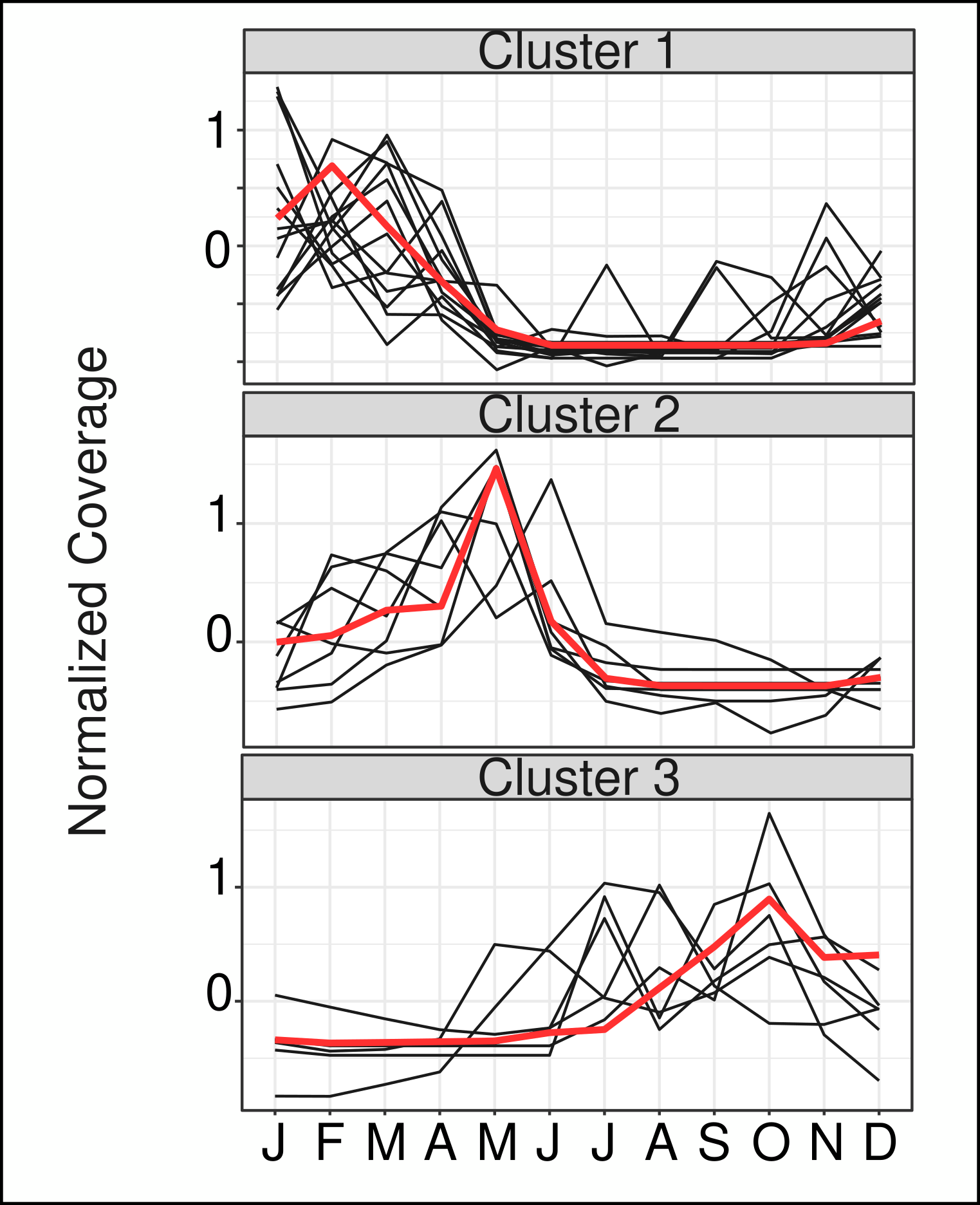

**Figure S5. 18S abundance clusters.** Clustering of 18S abundances was done using k-medoids clustering with the minimum Davies-Boudin index. Representative clusters are shown here with a red line denoting the medoid.

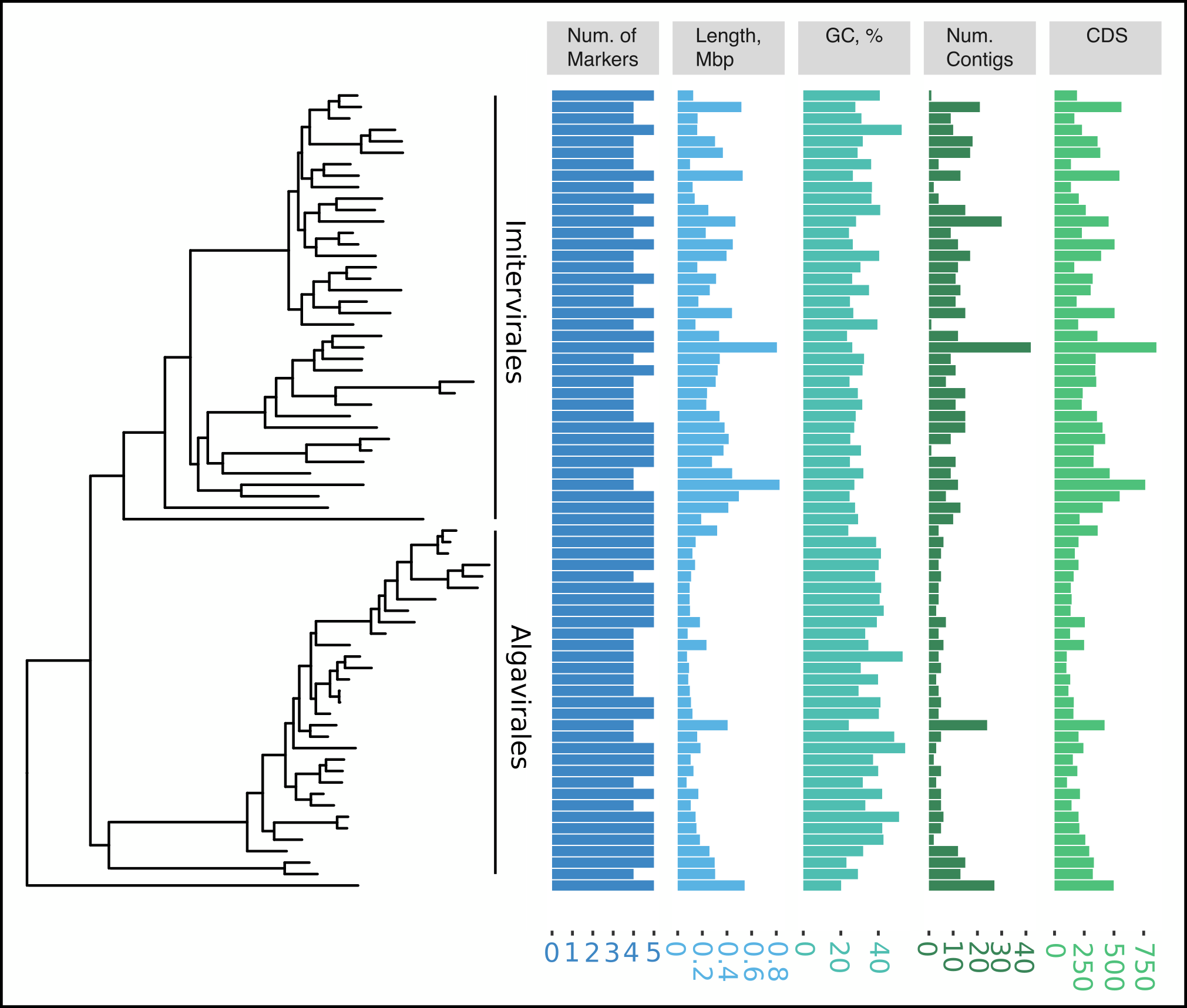

**Figure S6. Recovered NCLDV genome statistics.** The number of NCLDV marker genes, length in megabases, GC %, contig number, and coding sequences (CDS) are shown for each recovered NCLDV MAG. Phylogeny was generated as described in Figure 6.

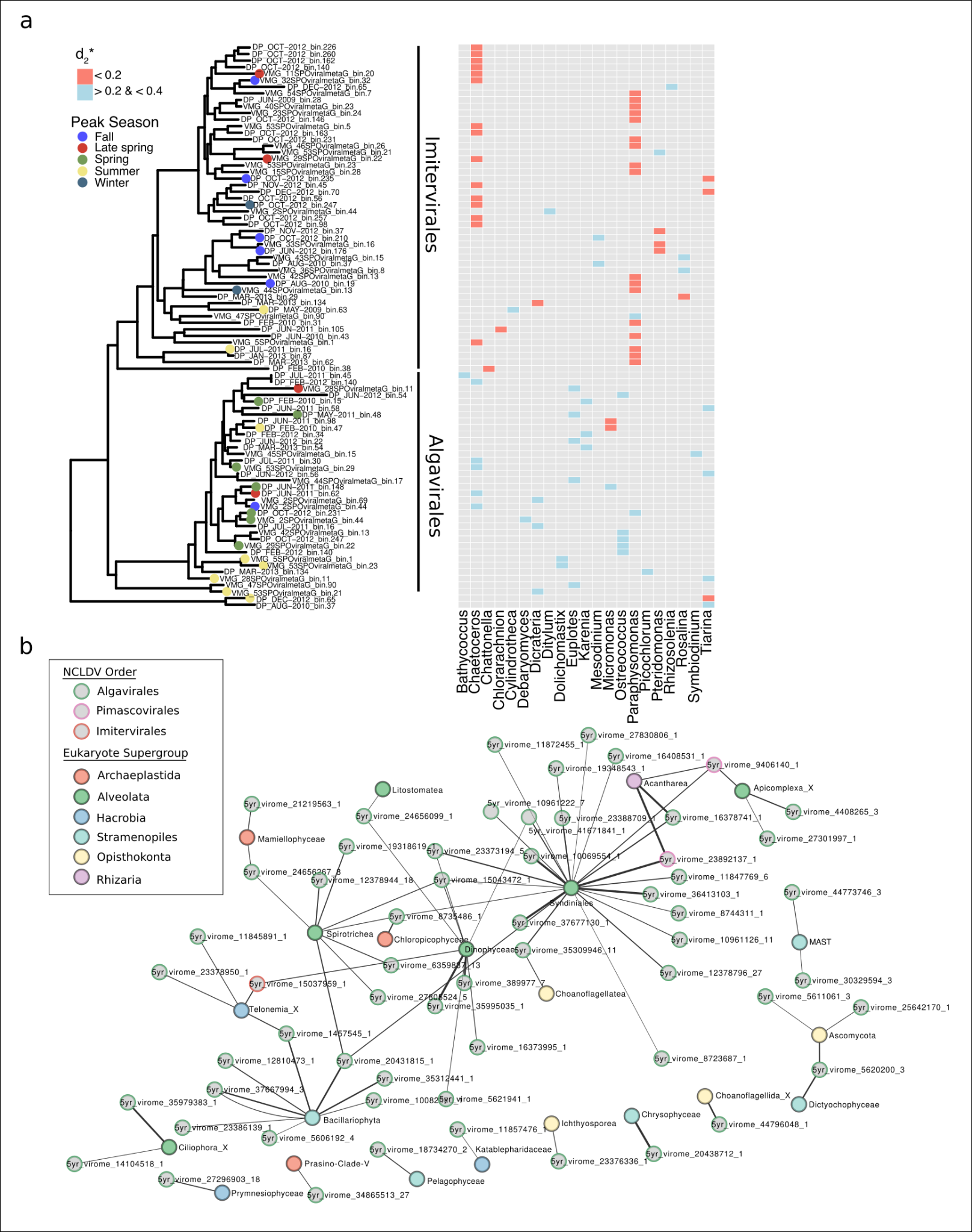

**Figure S7. Extended phylogenetic tree and network.** (a) The phylogenetic tree and host predictions from figure 7 are given leaf labels. (b) A network of host-virus interactions based on correlations between host 18s rRNA abundance and NCLDV phylotype abundance. Darker lines represent correlation strength.
